# Gonococcal invasion into epithelial cells depends on both cell polarity and ezrin

**DOI:** 10.1101/2021.04.29.441923

**Authors:** Qian Yu, Liang-Chun Wang, Daniel C. Stein, Wenxia Song

## Abstract

*Neisseria gonorrhoeae* (GC) establishes symptomatic infection in women from the cervix, lined with heterogeneous epithelial cells from non-polarized stratified at the ectocervix to polarized columnar at the endocervix. We have previously shown that GC differentially colonize and transmigrate across the ecto and endocervical epithelia. However, whether and how GC invade into heterogeneous cervical epithelial cells is unknown. This study examined GC entry of epithelial cells with various properties, using human cervical tissue explant and non-polarized/polarized epithelial cell line models. While adhering to non-polarized and polarized epithelial cells at similar levels, GC invaded into non-polarized more efficiently than polarized epithelial cells. The enhanced GC invasion in non-polarized epithelial cells was associated with increased ezrin phosphorylation, F-actin and ezrin recruitment to GC adherent sites, and the elongation of GC-associated microvilli. Inhibition of ezrin phosphorylation inhibited F-actin and ezrin recruitment and microvilli elongation, leading to a reduction in GC invasion. The reduced GC invasion in polarized epithelial cells was associated with non-muscle myosin II-mediated F-actin disassembly and microvilli ablation at GC adherence sites. Surprisingly, intraepithelial GC were only detected inside epithelial cells shed from the cervix, but neither in the ectocervix nor the endocervix, by immunofluorescence microscopy. We observed similar ezrin and F-actin recruitment in exfoliated cervical epithelial cells but not in those that remained in the ectocervical epithelium, as the luminal layer of ectocervical epithelial cells expressed ten-fold lower levels of ezrin than those beneath. However, GC inoculation induced F-actin reduction and myosin recruitment in the endocervix, similar to what was seen in polarized epithelial cells. Thus, polarized expression of ezrin at the apical surface of epithelial cells inhibits GC invasion, while non-polarized expression of ezrin promotes GC invasion by driving actin accumulation and microvilli elongation.

## Introduction

Gonorrhea, caused by a gram-negative bacterium *Neisseria gonorrhoeae* (GC), is the second most common sexually transmitted infection (STI) in the United States. It has recently become a public health crisis due to the emergence of antibiotic-resistant strains and a lack of vaccines [1–3]. A majority (50%~80%) of female gonococcal infections do not display any symptoms [4, 5]. GC can colonize the lower genital tract asymptomatically for a long time, and these asymptomatic infections can lead to severe complications [4]. When GC reach the upper genital tract, the infection can cause pelvic inflammatory disease (PID), a major cause of chronic adnominal pain, infertility, and predispose women to life-threatening ectopic pregnancy [6]. When entering the bloodstream, the infection can lead to disseminated gonococcal infection (DGI) [6, 7]. The asymptomatic nature of female infection allows GC to transmit silently and increases the risk of coinfections with HIV and other STIs [8].

The cervix is the gate from the unsterile lower female reproductive tract to the sterile upper tract [9] and a potential anatomic location that determines whether the outcome of GC infection is asymptomatic or symptomatic. The mucosal surface of the human cervix is characterized by heterogeneous epithelia, multiple-layered non-polarized stratified epithelial cells at the ectocervix and single-layered polarized columnar epithelial cells at the endocervix [10, 11]. The heterogeneous cervical epithelial cells build complicated physical and immune barriers against pathogens and also create a technical challenge for generating in vitro models that mimic the *in vivo* properties of the cervical surface to understand GC infection mechanism in females. We have established a human cervical tissue explant model [12] and shown that cervical epithelial cells in this explant model maintain most of the *in vivo* properties in culture and that GC infection in this explant model mimics what is observed in patients’ biopsies [13, 14].

GC establish infection by adhering to, invading into, and/or transmigrating across the epithelium [15–17]. GC express multiple adhesive molecules that can undergo phase and antigenic variation, such as pili, opacity-associated proteins (Opa), and lipooligosaccharide (LOS) [18–21]. These surface molecules mediate GC-epithelial interactions by binding to host cell receptors, establishing colonization. Pili initiate the interaction [22, 23], and Opa and LOS build intimate interactions of GC with the plasma membrane of epithelial cells [15, 17]. Such interactions induce epithelial cell signaling and cytoskeleton reorganization, which leads to microvilli elongation and membrane ruffling for GC entry into epithelial cells [24–27]. We have previously shown that GC interaction with polarized epithelial cells induces the disassemble of the apical junction, which allows GC transmigration across the epithelium [14]. GC-induced epithelial junction disassembly requires calcium-dependent activation of non-muscle myosin II (NMII), a part of the actomyosin supporting structure for the apical junction [14]. Using the human cervical tissue explant model, we have shown that GC differentially infect the various regions of the cervix when the different regions are equally exposed to the bacteria. GC preferentially colonize the ectocervical epithelium, using an integrin β1-dependent mechanism, and exclusively penetrate into the subepithelium of the endocervix by disrupting the apical junction using the NMII-dependent mechanism [13]. Pili are essential for GC colonization of all cervical regions. Expression of Opa proteins that bind carcinoembryonic antigen-related cell adhesion molecules (CEACAMs) as host receptors enhances colonization but reduces GC penetration [13]. However, whether and how GC invade into various cervical epithelial cells remains unknown.

Polarity is one of the properties that distinguish the ecto and endocervical epithelial cells. Ectocervical epithelial cells are flat and stratified with relatively smooth surfaces and low levels of cell polarity. In contrast, endocervical epithelial cells are tall and columnar with densely packed microvilli at the apical surface and high levels of cell polarity [10, 28]. Polarized columnar endocervical epithelial cells form the apical junction complexes with neighboring cells, which hold columnar epithelial cells together into a monolayer, seal spaces between cells to build a physical barrier, and facilitate the polarized distribution of surface molecules [29]. The actin cytoskeleton in ectocervical epithelial cells shows no polarity, distributing evenly at the cell periphery, supporting the cell membrane. In contrast, endocervical epithelial cells exhibit strong polarity of the actin cytoskeleton, concentrating under the apical membrane and supporting the apical junction and dense microvilli [30]. Actin-mediated morphological changes in both non-polarized and polarized epithelial cells, including microvilli formation, have been shown to require ezrin [31, 32]. Upon phosphorylation, ezrin interacts with both F-actin and membrane proteins, linking the actin cytoskeleton to the plasma membrane of epithelial cells [33]. Ezrin has also been suggested to play a role in microbial infection of epithelial cells, including GC [34, 35], *Neisseria meningitis* [36, 37], *Chlamydia trachomatis* [38], *Group A Streptococcus* [39], *Enteropathogenic Escherichia coli* [40], and *Pseudomonas aeruginosa* [41]. GC interaction with non-polarized epithelial cells, including primary cervical epithelial cells, induces ezrin recruitment [25, 35]. Whether differences in epithelial cell polarity and morphology and the underlying actin organization impact GC infection at various cervical regions is unclear.

This study examined the effect of polarity, morphology, and actin organization in ecto and endocervical epithelial cells on GC invasion and the underlying mechanism, using a non-polarized and polarized epithelial cell line model generated from the same cell line and the human cervical tissue explant model. We found that the polarity of epithelial cells and the expression of ezrin are two critical factors determining the efficiency of GC invasion into epithelial cells. We further investigated the mechanisms by which the polarity of cervical epithelial cells and the level of ezrin expression regulate GC invasion.

## Results

### GC preferentially invade non-polarized epithelial cells

To determine if cell polarity alone affects GC infectivity, we established non-polarized and polarized epithelial cells on transwells using human colonic epithelial cell line T84. After comparing multiple types of human epithelial cells, including endometrial epithelial cells HEC-1-B, primary cervical epithelial cells [25], and cervical epithelial cells immortalized by HPV [42], we chose T84 as it was the only cell line that can be polarized to the level and exhibit the columnar morphology and the barrier function, similar to endocervical epithelial cells *in vivo* (S1 Fig and S2 Fig). T84 cells also express CEACAMs, the host receptors for most variants of GC Opa proteins [43]. The two-day culture of T84 cells on transwells was modeled as non-polarized cells, as monolayers had low transepithelial resistance (TEER) (<500 Ω), were permeable to the CellMask dye (S1 Fig), and showed flat morphology and random distribution of F-actin and the junction proteins E-cadherin and ZO-1 (S2 Fig). After culture on transwells for ~10 days, T84 cells became polarized, exhibiting > 2000 Ω TEER, the barrier function against the dye (S1 Fig), and apical polarization of F-actin, E-cadherin, and ZO-1 staining (S2 Fig).

Using the cell line model, we compared the effect of epithelial morphology and polarity on the adhesion and invasion of MS11 GC that was piliated and expressed phase variable Opa (Pil+Opa+) or no Opa (Pil+ΔOpa) using the gentamicin resistance assay [26, 44]. After 3-h inoculation, both Pil+Opa+ and Pil+ΔOpa GC adhered to non-polarized and polarized epithelial cells at similar levels (Fig. 1A). In contrast, significantly more gentamicin-resistant Pil+Opa+ and Pil+ΔOpa GC were recovered in the non-polarized than the polarized epithelial cells after 6-h inoculation (Fig. 1B). Furthermore, Pil+Opa+ GC exhibited significantly higher levels of adherence and invasion in both non-polarized and polarized T84 cells than Pil+ΔOpa GC (Fig. 1A and 1B). To confirm the enhanced invasion of Pil+Opa+ GC in the non-polarized epithelial cells, we used transmission electron microscopy (TEM). We quantified the percentage of intracellular GC over the total epithelial cell-associated GC in individual images (Fig. 1C and 1D). We observed ~16% Pil+Opa+ and 7% Pil+ΔOpa GC inside non-polarized cells, but no intracellular GC were found inside polarized epithelial cells (Fig. 1D). These results suggest that GC preferentially invade into non-polarized epithelial cells, and Opa expression enhances both GC adherence and invasion into non-polarized and polarized epithelial cells.

**Fig. 1.**
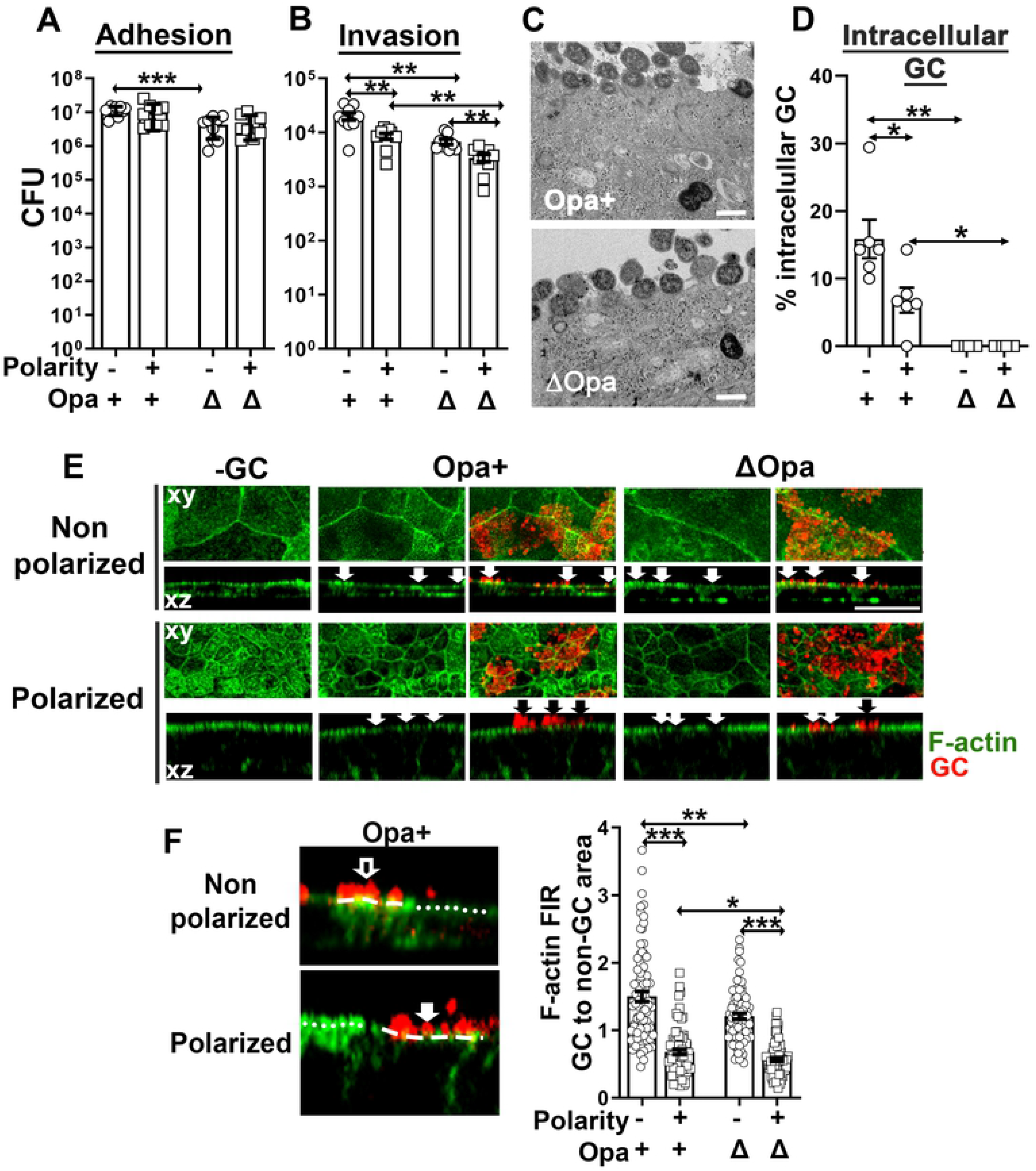
The polarity of epithelial cells regulates GC infection potentially by inducing distinct actin cytoskeleton responses. **(A, B)** T84 cells cultured 2 or 10 days were inoculated with Pil+Opa+ or ΔOpa GC (MOI~10) from the top chamber. Adherent GC (±SEM) were quantified by culturing the lysates of infected epithelial cells 3 h post-incubation **(A)**. Invaded GC (±SEM) were quantified by gentamicin resistance assay 6 h post-inoculation **(B)**. Data points represent individual transwells. **(C, D)** Cells were incubated with Pil+Opa+ or Pil+ΔOpa GC (MOI~50) from the top chamber for 6 h, fixed, and processed for transmitted electron microscopy (TEM). Representative images of non-polarized T84 cells with intracellular GC are shown **(C)**. The level of invasion was quantified by percentages of the intracellular GC among the total epithelial-associated GC **(D)**. Data points represent individual images. **(E, F)** Cells were treated as A, B, fixed 6 h post-inoculation, stained for F-actin and GC, and analyzed using 3D-CFM. Representative xy images of the top surface and xz images crossing the top and the bottom surfaces are shown. Arrows indicate the location of GC **(E)**. The redistribution of F-actin was quantified by the mean fluorescence intensity ratio (FIR) (±SEM) of F-actin underneath individual GC microcolonies relative to the adjacent no GC surface area **(F)**. Data points represent individual GC microcolonies. Scale bar, 20 μm. n=3, \**p*<0.05; \*\**p*< 0.01; \*\*\**p*<0.001.

### GC inoculation induces distinct F-actin reorganization in non-polarized and polarized epithelial cells

The actin cytoskeleton is required for GC invasion [26, 45], and non-polarized and polarized epithelial cells organize actin differently (S2 Fig). We postulated that the difference in actin organization in non-polarized and polarized epithelial cells is one of the underlying causes of differential GC invasion efficiencies. To test this hypothesis, we inoculated non-polarized and polarized T84 cells with Pil+Opa+ and Pil+ΔOpa GC for 6 h, fixed, stained for F-actin with phalloidin and GC with an antibody, and analyzed by three-dimensional confocal fluorescence microscopy (3D-CFM) to examine cells from the top (xy view, Fig. 1E top panels) and the side (xz view, Fig. 1E, bottom panels) of the epithelium. We observed different actin reorganization beneath GC adherent sites in non-polarized and polarized epithelial cells (Fig. 1E, arrows). We quantified the actin reorganization by the mean fluorescent intensity ratio (FIR) of F-actin staining underneath GC microcolonies relative to the adjacent no GC surface area. The FIRs of F-actin staining underneath both Pil+Opa+ and Pil+ΔOpa microcolonies were higher than 1 in non-polarized epithelial cells but lower than 1 in polarized epithelial cells (Fig. 1E and 1F), indicating the enrichment and reduction of F-actin at GC adherent sites in non-polarized and polarized epithelial cells, respectively. When comparing the two GC strains, the F-actin FIRs underneath Pil+Opa+ GC were significantly higher than those underneath Pil+ΔOpa GC in both non-polarized and polarized epithelial cells (Fig. 1F). These data indicate that GC induce opposite actin reorganization in non-polarized (F-actin enrichment) and polarized epithelial cells (F-actin reduction) at adherent sites. Opa expression enhances the F-actin enrichment in non-polarized epithelial cells and alleviates the F-actin reduction in polarized epithelial cells.

### GC inoculation activates and recruits ezrin to adherent sites in non-polarized epithelial cells, facilitating F-actin enrichment and GC invasion

Ezrin functions as a linker between the actin cytoskeleton and the plasma membrane, and its linker function is activated through phosphorylation of threonine 567 [33]. We determined whether GC induced differential actin reorganization in non-polarized and polarized epithelial cells through ezrin by analyzing ezrin cellular distribution and phosphorylation in response to GC inoculation. We inoculated non-polarized and polarized T84 cells with Pil+Opa+ and Pil+ΔOpa GC for 6 h. Cells were fixed, stained for ezrin and GC, and analyzed by 3D-CFM. We evaluated the redistribution of ezrin staining underneath GC adherent sites relative to the adjacent no GC surface area by FIR. In the absence of GC, the ezrin staining was distributed randomly in the cytoplasm and periphery of non-polarized epithelial cells, but concentrated at the apical surface in polarized epithelial cells (Fig. 2A), similar to the distribution of F-actin (Fig. 1E). Inoculation of Pil+Opa+ or Pil+ΔOpa GC both induced ezrin enrichment at adherent sites of non-polarized epithelial cells (Fig. 2A, arrows), leading to ezrin FIRs > 1 (Fig. 2B). Furthermore, the ezrin FIR in Pil+Opa+ GC-inoculated non-polarized epithelial cells was significantly higher than in Pil+ΔOpa GC-inoculated cells (Fig. 2B). In contrast, we observed no change or a reduction in ezrin staining at Pil+Opa+ and Pil+ΔOpa GC adherent sites, respectively, at the apical surface of polarized epithelial cells (Fig. 2A, arrows). Consequently, their ezrin FIRs were close to or smaller than one, respectively, while both were significantly lower than those in non-polarized epithelial cells (Fig. 2B). Thus, similar to GC-induced actin reorganization, GC inoculation causes ezrin recruitment and disassociation at adherent sites in non-polarized and polarized epithelial cells, respectively. Opa expression enhances the ezrin recruitment in non-polarized epithelial cells but alleviates the ezrin disassociation in polarized epithelial cells.

**Fig. 2.**
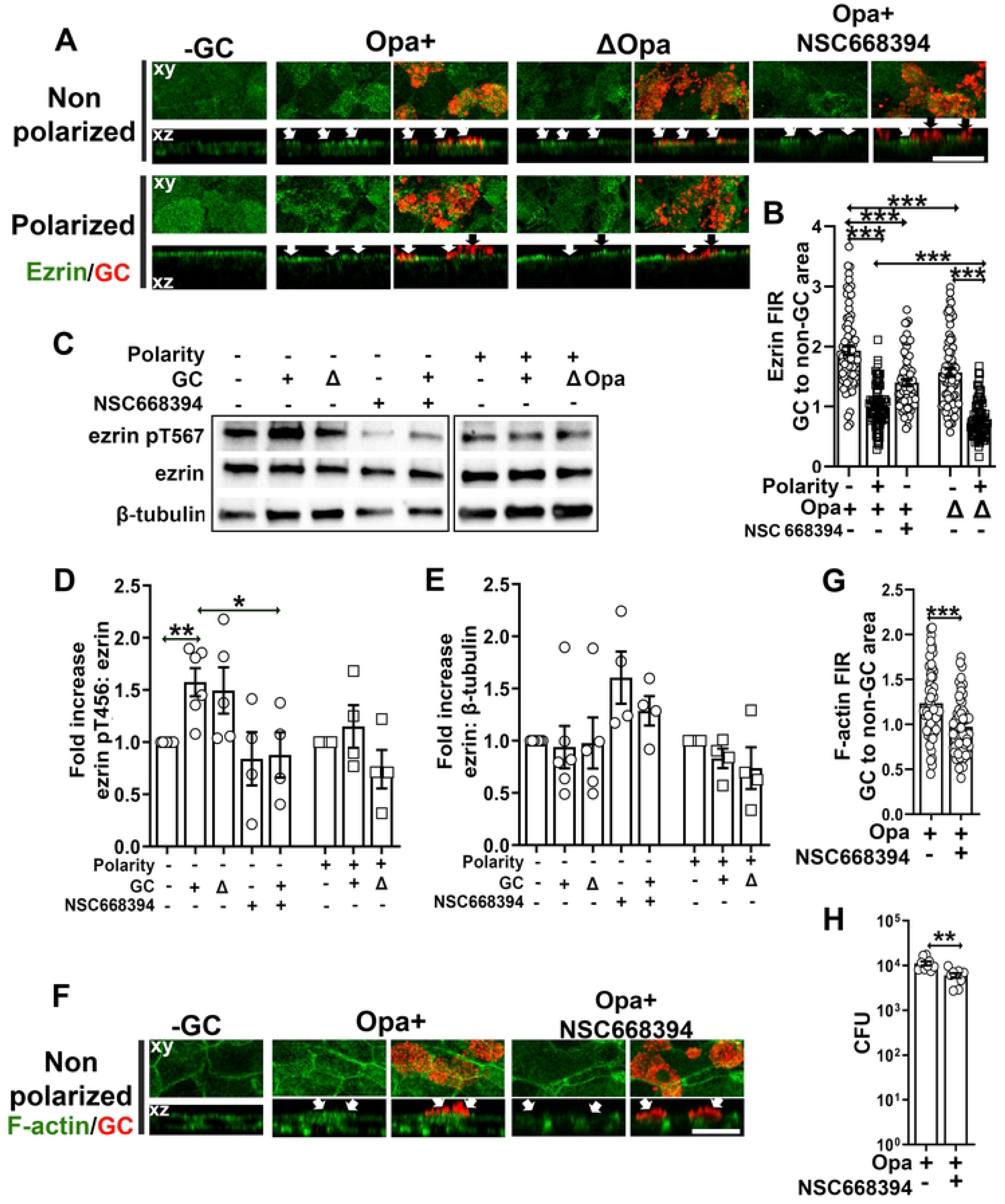
GC induce differential activation and redistribution of ezrin in non-polarized and polarized epithelial cells for actin reorganization and GC invasion. T84 cell 2-day and 10-day cultures were pretreated with or without the ezrin activation inhibitor NSC668394 (20 μM) for 1 h and apically inoculated with Pil+Opa+ or ΔOpa GC (MOI~10) for 6 h with or without the inhibitor. **(A, B, F, G)** Cells were fixed, permeabilized, stained for ezrin **(A)**, F-actin **(F)**, and GC, and analyzed using 3D-CFM. Shown are representative xy and xz images **(A, F)**. The levels of ezrin **(B)** and F-actin **(G)** redistribution were quantified by FIR (±SEM) underneath GC microcolonies relatively to the adjacent no GC surface area. Data points represent individual GC microcolonies. **(C-E)** Cells were lysed and analyzed by western blot. Shown are representative blots **(C)**. The average fold of increase in the ezrin pT567 to ezrin ratio **(D)** and the ezrin to β-tubulin ratio **(E)** (±SEM) was quantified by NIH ImageJ. Data points represent individual transwells. **(H)** Invaded GC (±SEM) were quantified by gentamicin resistance assay 6 h post-inoculation with or without NSC668394 treatment. Data points represent individual transwells. Scale bar, 20 μm. n=2~3, \**p*<0.05; \*\**p*< 0.01; \*\*\**p*<0.001.

GC-induced ezrin redistribution suggests a role for GC in regulating ezrin activation. To test this, we determined the levels of ezrin phosphorylation at T567 (pT567) using western blotting 6 h post-incubation (Fig. 2C). Using the density ratio of ezrin pT567 and total ezrin, we found a significant increase in ezrin phosphorylation in GC inoculated non-polarized epithelial cells compared to no GC controls (Fig. 2C and 2D). However, GC inoculation did not significantly change the phosphorylation level of ezrin in polarized epithelial cells. We did not detect any significant difference in ezrin phosphorylation levels between Pil+Opa+ and Pil+ΔOpa GC-inoculated epithelial cells, no matter if they were polarized or not. There was also no significant difference in the density ratio of ezrin to β-tubulin in response to GC inoculation in both non-polarized and polarized epithelial cells (Fig. 2C and 2E). Thus, GC inoculation induces ezrin activation exclusively in non-polarized epithelial cells. The above data suggest that GC induced the activation and recruitment of ezrin to adherent sites in non-polarized epithelial cells but not and even reduced ezrin at adherent sites in polarized epithelial cells.

The activation and recruitment of ezrin to GC adherent sites suggest a role for ezrin in GC-induced actin reorganization and GC infection. We examined this hypothesis utilizing an ezrin inhibitor NSC668394, which blocks ezrin phosphorylation at threonine 567 [46]. We determined the effect of the inhibitor on ezrin and actin reorganization by immunofluorescence microscopy and FIR as described above and on GC invasion by the gentamicin resistant assay. The ezrin inhibitor significantly reduced the FIRs of ezrin underneath GC adherent sites relative to the no GC area (Fig. 2A and 2B) and ezrin phosphorylation without affecting the protein level of ezrin (Fig. 2C-2E), indicating effective inhibition of ezrin activation. Importantly, the inhibitor treatment significantly reduced the FIRs of F-actin underneath GC adherent sites relative to the no-GC area (Fig. 2F and 2G) as well as the invasion level of Pil+Opa+ GC, compared to no inhibitor controls (Fig. 2H). However, the ezrin inhibitor had no significant effect on GC growth (S3 Fig). These data indicate that the activation and recruitment of ezrin to GC adherent sites in non-polarized epithelial cells directly leads to F-actin accumulation, facilitating GC invasion.

### GC induce the activation and recruitment of NMII to GC adherent sites in polarized epithelial cells exclusively, causing actin depolymerization

We have previously shown that GC inoculation induces NMII activation and recruitment to GC adherent sites in polarized epithelial cells in a calcium-dependent manner, leading to apical junction disassembly and gonococcal transmigration across the epithelium [14]. Here, we addressed whether NMII, an actin motor, was involved in the differential F-actin remodeling in non-polarized and polarized epithelial cells. We first determined whether GC could induce NMII activation and redistribution in non-polarized epithelial cells, similarly to what we observed in polarized epithelial cells, using immunofluorescence microscopy. In the absence of GC, phosphorylated NMII light chain (pMLC) staining was distributed randomly at the cell periphery and the cell-cell contact in non-polarized epithelial cells but concentrated at the apical part of polarized epithelial cells (Fig. 3A). Inoculation with Pil+Opa+ or Pil+ ΔOpa GC for 6 h did not significantly change the distribution of pMLC in non-polarized epithelial cells, and the FIRs of pMLC staining at GC adherent sites compared to no GC surface area was near 1 (Fig. 3A, arrows, and 3B). This was in sharp contrast with what occurred in polarized epithelial cells – pMLC concentrated at GC adherent sites at the apical surfaces, resulting in the FIRs of pMLC at GC adherent sites relative to no GC surface area higher than 1 (Fig. 3A, arrows, and 3B). Pil+ΔOpa-infected polarized epithelial cells exhibited significantly higher pMLC FIRs (~ 1.9) than Pil+Opa+ GC-infected polarized epithelial cells (~ 1.6) (Fig. 3B). These data suggest that GC inoculation only induced the activation and recruitment of NMII in the polarized but not non-polarized epithelial cells, and Opa expression suppresses such effects.

**Fig. 3.**
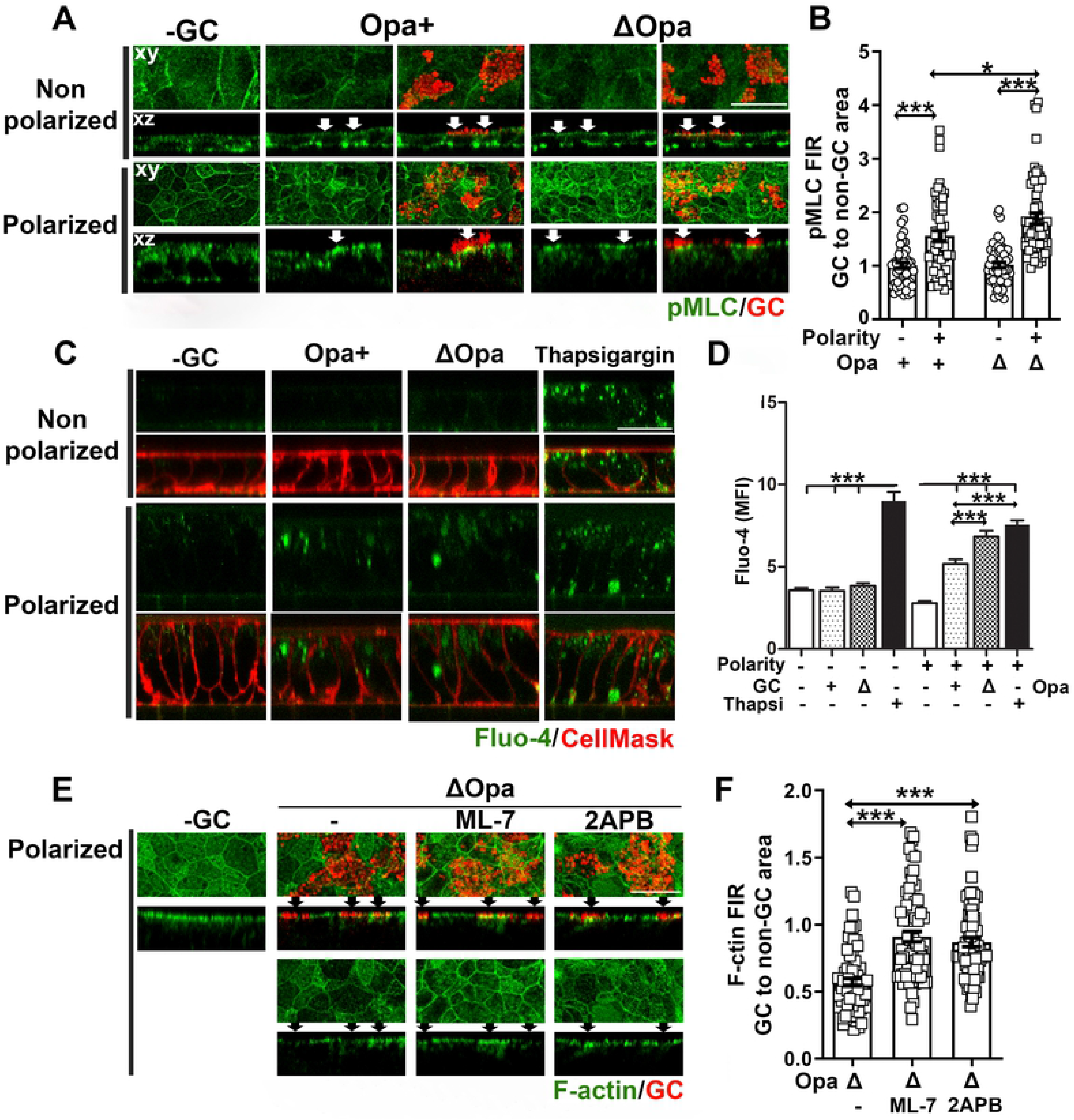
NMII activation and Ca^2+^ elevation are required for GC-induced F-actin reduction at adherent sites of polarized epithelial cells. **(A, B)** Non-polarized and polarized T84 cells were inoculated with Pil+Opa+ or ΔOpa GC from the top chamber for 6 h (MOI~10). Cells were fixed and stained for phosphorylated myosin light chain (pMLC) and GC and analyzed using 3D-CFM. Shown are representative xy images of the top surface, xz images crossing both the top and the bottom surface **(A),** and the FIR (±SEM) of pMLC staining underneath individual GC microcolonies relative to the adjacent no GC surface area **(B)**. **(C, D)** Non-polarized and polarized T84 cells were incubated from the top chamber with or without GC Pil+Opa+ or Pil+ΔOpa in the presence or absence of thapsigargin (10 μM) for 4 h. Cells were incubated with the Ca^2+^ indicator Fluo4 and the membrane dye CellMask and analyzed using 3D-CFM. Shown are representative xz images **(C)** and the mean fluorescence intensity (MFI) (±SEM) of Fluo-4 in the cytoplasmic region of individual epithelial cells **(D)**. **(E, F)** Polarized T84 cells were pre-treated with or without ML-7 (10 μM) or 2APB (10 μM) and apically inoculated with Pil+ΔOpa GC in the presence or absence of inhibitors for 6 h (MOI~10). Cells were fixed and stained for F-actin using phalloidin and GC using antibody and analyzed using 3D-CFM. Shown are representative xy and xz images **(E)** and the FIR (±SEM) of phalloidin staining underneath individual GC microcolonies relative to the adjacent no GC surface area **(F)**. Data points in (B) and (F) represent individual GC microcolonies. Scale bar, 20 μm. n=3, \**p*<0.05; \*\*\**p*<0.001.

We next determined whether the absence of the NMII activation and recruitment in non-polarized epithelial cells is due to a lack of Ca^2+^ flux response exhibited by polarized epithelial cells [14], using Fluo-4, a calcium dye, and 3D-CFM. Thapsigargin treatment, which elevates intracellular Ca^2+^ [47], served as a positive control. Consistent with previously published studies [14], GC inoculation increased the intracellular Ca^2+^ level of polarized epithelial cells, with the Ca^2+^ level in Pil+ΔOpa GC-infected cells significantly higher than Pil+Opa+ GC-infected cells (Fig. 3C and 3D). However, GC inoculation did not change Ca^2+^ levels in non-polarized epithelial cells (Fig. 3C and 3D). This result suggests that GC inoculation only elevates the cytoplasmic Ca^2+^ levels in polarized epithelial cells but not in non-polarized epithelial cells.

To determine whether GC-induced activation of NMII and Ca^2+^ flux contributed to GC-induced actin reorganization in polarized epithelial cells, we utilized an MLC kinase (MLCK) inhibitor, ML-7 [48], and a Ca^2+^ flux inhibitor, 2APB [49], which have been shown to effectively inhibit GC-induced activation of NMII and Ca^2+^ flux [14]. We evaluated their effects on the F-actin reduction at adherent sites of Pil+Opa+ GC in polarized epithelial cells, using 3D-CFM and the FIR of F-actin at GC adherent sites relative to the no GC surface area (Fig. 3E and 3F). Both the MLCK and Ca^2+^ inhibitors restored the level of F-actin at GC adherent sites and significantly increased the F-actin FIR from ~0.6 to 0.9 (Fig. 3E and 3F). These data together suggest that GC reduce F-actin at adherent sites in polarized epithelial cells by activating Ca^2+^ flux and NMII locally.

### GC differentially remodel microvilli of non-polarized and polarized epithelial cells depending on ezrin- and NMII, respectively

Membrane ruffling and microvilli elongation have been observed in inoculated primary cervical and endometrial epithelial cells [25, 50] and cell lines [24, 26], as well as patient biopsies [51]. Such morphological changes have been implicated for GC invasion. We determined whether the differential actin reorganization we observed causes different morphological changes in non-polarized and polarized epithelial cells using transmitted electron microscopy (TEM). Without GC inoculation, there are short irregular membrane protrusions randomly distributing on the top surface of non-polarized epithelial cells (Fig. 4A, top panels). With GC inoculation, the epithelial membrane contacting Pil+Opa+ or Pil+ΔOpa GC showed elongated microvilli wrapping around bacteria (Fig. 4A, arrows). We evaluated these morphological changes by measuring the percentage of GC contacting elongated microvilli directly in individual TEM images. We found that ~47% Pil+Opa+ GC and ~33% of Pil+ΔOpa GC with elongated microvilli (Fig. 4 B). Differing from non-polarized epithelial cells, polarized cells have densely packed vertical microvilli at the apical surface in the absence of GC (Fig. 4A, top panels). Post inoculation, the microvilli contacting Pil+Opa+ GC became shorter and bent (Fig. 4C open arrows). The epithelial membrane contacting Pil+ΔOpa GC lost most microvilli, allowing close interactions of GC with the apical membrane of polarized epithelial cells (Fig. 4C, open arrowheads). Quantitative analysis of TEM images showed ~63% Pil+ΔOpa but only ~13% Pil+Opa+ GC causing microvilli disappearance (Fig. 4D). These results indicate that GC induce microvilli elongation in non-polarized epithelial cells but destroy microvilli at the contacting membrane of polarized epithelial cells. Opa expression promotes microvilli elongation in non-polarized epithelial cells while protecting microvilli from damage in polarized epithelial cells.

**Fig. 4.**
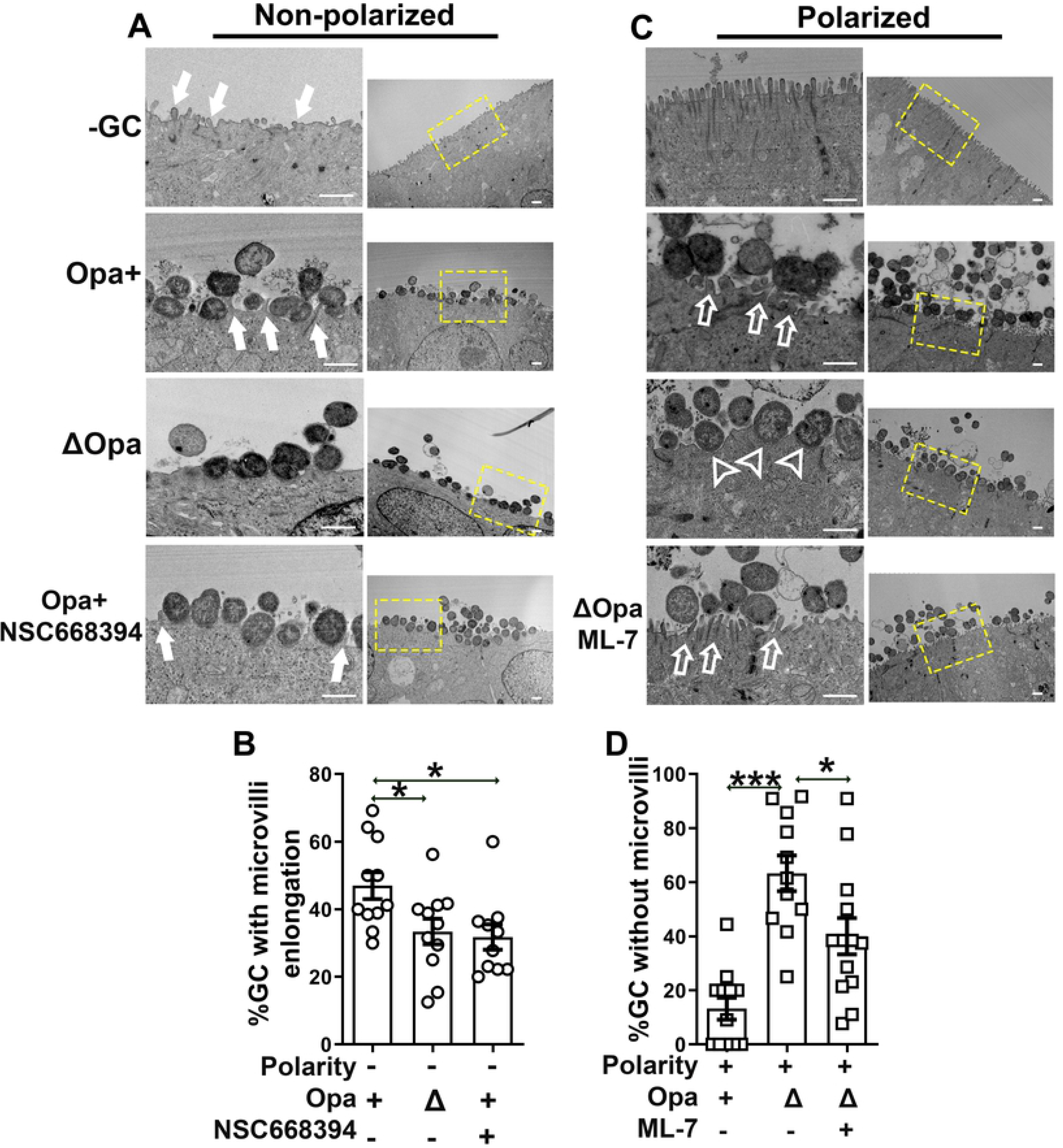
GC differentially remodel microvilli of non-polarized and polarized epithelial cells depending on ezrin and NMII, respectively. Non-polarized and polarized T84 cells were pre-treated with or without NSC668394 (20 μM) or ML-7 (10 μM) and incubated with Pil+Opa+ or Pil+ΔOpa GC (MOI~50) in the apical chamber for 6 h with or without the inhibitors. Cells were fixed and processed for transmitted electron microscopy (TEM). **(A, C)** Shown are representative images with high (left panels) and low (right panels) magnifications. Yellow dash line rectangles highlight the focused area. Solid arrows, GC-associated elongated microvilli. Open arrows, GC-induced microvilli bending. Open arrowheads, GC-induced ablation of microvilli. **(B, D)** Membrane morphological changes were quantified by the percentage of GC associating with elongated microvilli in non-polarized epithelial cells **(B)** and the percentage of GC associating with the plasma membrane of polarized epithelial cells that lost microvilli **(D)**. Data points represent individual randomly acquired TEM images. Scale bar, 1 μM. n=2-3. \**p*<0.05; \*\*\**p*<0.001.

To answer whether GC-induced differential actin reorganization was responsible for the epithelial morphological changes, we examined the effect of the ezrin activation inhibitor NSC668394 and the MLCK inhibitor ML-7 on GC-associated microvilli. Treatment of NSC668394 significantly reduced the percentage of Pil+Opa+ GC associating with elongated microvilli from ~47% to ~32% in non-polarized epithelial cells (Fig. 4A, arrows, and 4B). Conversely, treatment of ML-7 significantly reduced the percentage of GC associating with the epithelial membrane that lost microvilli from ~63% to ~40% in polarized epithelial cells (Fig. 4C, open arrows, and 4D). These data indicate that GC adherence induces microvilli elongation locally in non-polarized epithelial cells by activating ezrin to promote invasion, but locally destroys microvilli at the apical surface of polarized epithelial cells by activating NMII and NMII-driven actin depolymerization to facilitate GC-epithelial contact. Opa expression promotes ezrin- and actin-driven microvilli elongation in non-polarized epithelial cells but inhibits NMII-driven ablation of microvilli in polarized epithelial cells.

### No significant GC were detected inside the ectocervical and endocervical epithelia

We evaluated whether GC can invade into the epithelial cells in the human cervix. Human cervical tissue explants were incubated with Pil+Opa+ and Pil+ΔOpa GC (MOI~10) for 24 h, washed at 6 and 12 h post-inoculation to remove unassociated bacteria, and cryopreserved. Tissue sections were stained for GC, F-actin and DNA, and analyzed using CFM as previously described [13]. GC staining was predominantly detected at the luminal surface of both stratified ectocervical and columnar endocervical epithelial cells, occasionally in the subepithelium of the endocervix, but hardly any within the epithelium (Fig. 5A). We evaluated GC distribution in cervical tissue explants by measuring the percentage of GC fluorescence intensity (% GC FI) at the luminal surface, within the cervical epithelium, and at the subepithelium. Nearly 100% of GC staining was at the luminal surface of the ectocervical epithelium. No significant GC staining was detected within the ectocervical epithelium and at the subepithelium, no matter which strain of GC was used (Fig. 5B). While significant percentages of GC staining were detected at the endocervical subepithelium, the percentage of GC staining within the endocervical epithelium remained very low (Fig. 5B). Consistent with our previously published data [13], the percentage of Pil+ΔOpa GC staining at the endocervical subepithelium was significantly higher than that of Pil+Opa+ GC (Fig. 5, B). While distinguishing intracellular and extracellular GC staining in tissue explants was technically difficult, we noticed GC staining inside epithelial cells shedding from cervical tissues (Fig. 5D, open arrows, and S4A Fig, open arrows). However, shedding cervical cells were rarely caught as they were likely disassociated from tissues. Consequently, quantification became impossible.

**Fig. 5.**
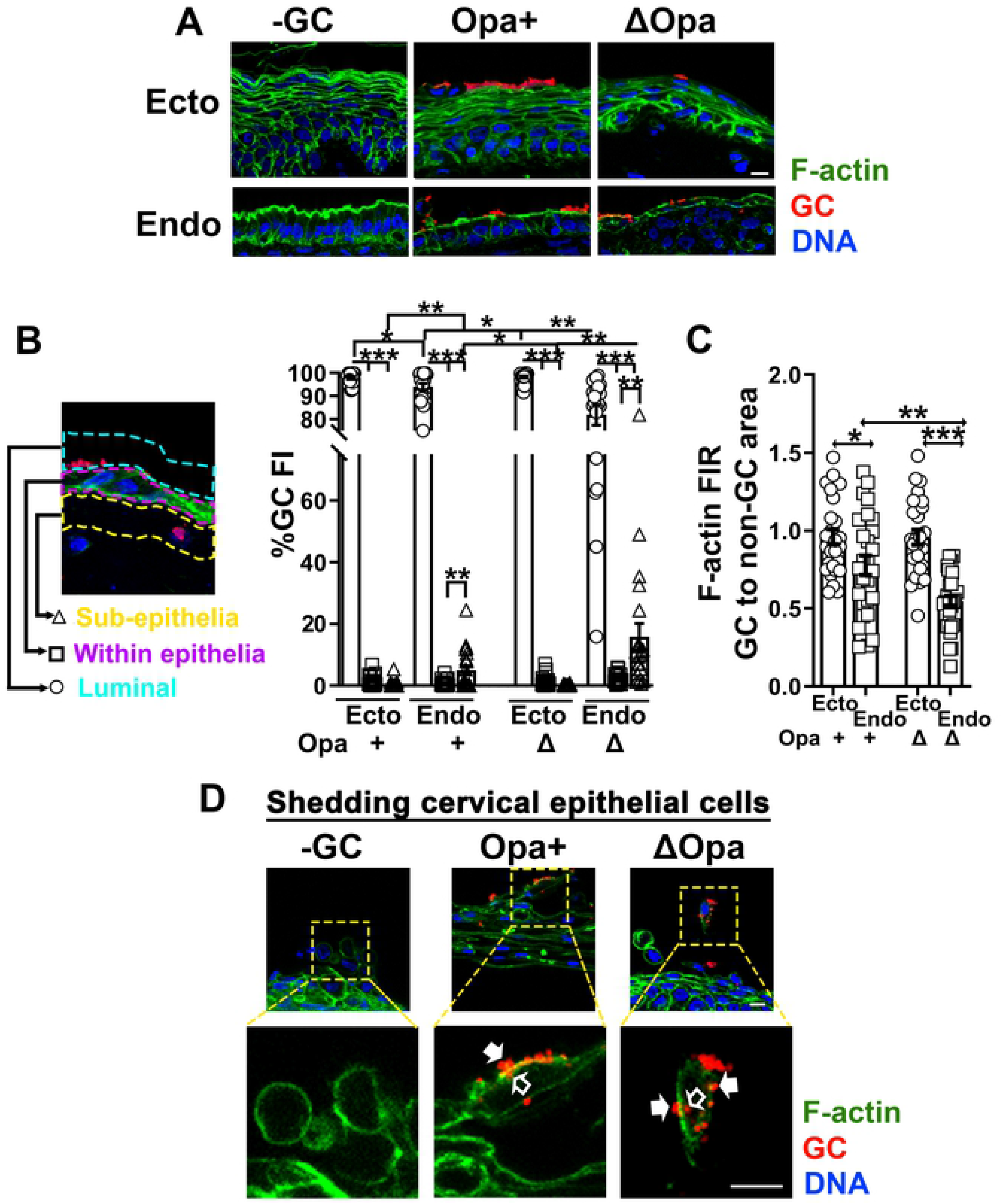
The level of intraepithelial GC is much lower than those on the luminal surface and/or in the subepithelia of the ecto- and endocervix. Human cervical tissue explants were incubated with Pil+Opa+ and Pil+ΔOpa (MOI~10) for 24 h, with unassociated GC washed off at 6 and 12 h. Tissue explants were fixed, stained for F-actin, GC and DNA, and analyzed using CFM. **(A)** Shown are representative images of the ecto and endocervix. Scale bar, 10 μm. **(B)** The distribution of GC in cervical tissues was quantified by the percentage of GC fluorescent intensity (% GC FI) detected at the luminal surface (blue dash lines), within the epithelium (magenta dash lines), and at the subepithelium (yellow dash lines), relatively to the total GC FI in the entire area. Data points represent individual randomly acquired images. **(C)** The redistribution of F-actin was quantified by the FIR (±SEM) of phalloidin staining underneath individual GC microcolonies relative to the adjacent no GC luminal area. Data points represent individual GC microcolonies. **(D)** Shown are representative images of shedding cervical epithelial cells stained for GC, F-actin, and DNA. Open arrows, intracellular GC. Arrows, surface GC microcolonies with F-actin accumulation. Scale bar, 10 μm. n=3. \**p*<0.05; \*\**p*< 0.01; \*\*\**p*<0.001.

As distinct actin reorganization leads to different GC invasion levels in non-polarized and polarized epithelial cells, we examined the distribution of F-actin, stained by phalloidin, in cervical tissues in relation to GC locations, using the FIR of phalloidin staining at GC adherent sites relatively to an adjacent luminal area without GC. Contrary to the results from the non-polarized epithelial cell line model showed above, the FIR of F-actin remained at ~1 in stratified non-polarized ectocervical epithelial cells, indicating no F-actin accumulation at GC adherent sites (Fig. 5A and 5C). We observed F-actin accumulation at GC adherent sites of shedding cervical epithelial cells (Fig. 5D, arrows, and S4B Fig, arrows). Similar to what we observed in polarized T84 cells, the F-actin FIR in GC colonized columnar polarized endocervical epithelial cells was reduced below 1 (Fig. 5A and 5C), showing F-actin reduction at GC adherent sites. The F-actin FIR in Pil+ΔOpa GC-colonized was significantly lower than that in Pil+Opa+ GC-colonized endocervical epithelial cells (Fig. 5C).

These data suggest that the number of GC invading into cervical epithelial cells is less than the numbers of GC colonizing on the luminal surfaces of the ecto- and endocervix and penetrating the subepithelium of the endocervix, even though GC may readily invade into epithelial cells shedding off the cervix. The low levels of intraepithelial GC in the cervix may be associated with GC failure to induce F-actin accumulation at adherent sites.

### GC fail to induce ezrin and NMII reorganization in ectocervical epithelial cells

To understand our surprising findings of a lack of intraepithelial GC and F-actin accumulation at GC adherent sites in the ectocervix, we examined ezrin and NMII responses of ectocervical epithelial cells to GC inoculation, using the tissue explant model and immunofluorescence microscopy. In the absence of GC inoculation, we detected ezrin and pMLC staining in the subluminal layers but very little of both in the luminal layer of the ectocervical epithelium (Fig. 6A and 6C). We compared the staining levels of ezrin or pMLC between the luminal and immediate subluminal layers of the ectocervical epithelium. The MFI of both ezrin (Fig. 6B) and pMLC (Fig. 6D) in the luminal layer of epithelial cells were ~10 times less than that in the subluminal layer. Furthermore, GC inoculation did not increase ezrin and pMLC staining at GC adherent sites of the luminal ectocervical epithelial cells (Fig. 6A-D, arrows). However, we detected the accumulation of ezrin staining underneath microcolonies of both Pil+Opa+ and Pil+ΔOpa GC in shedding cervical epithelial cells (Fig. 6E, arrows). These data indicate that GC fail to induce the recruitment of both ezrin and NMII to adherent sites in ectocervical epithelial cells, likely due to their low expression in the luminal layer.

**Fig. 6.**
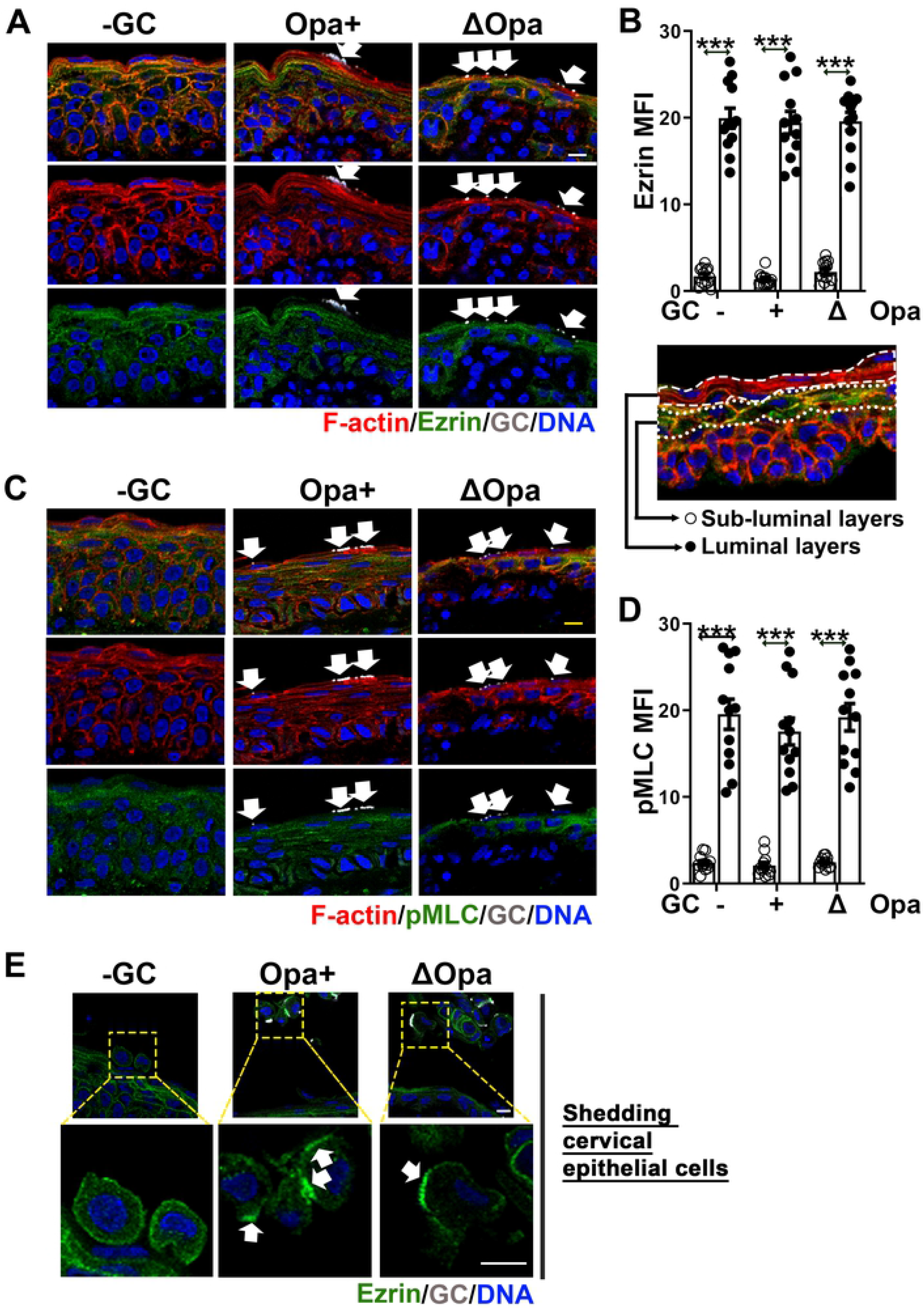
GC fail to induce ezrin and NMII reorganization in ectocervical epithelial cells. Human cervical tissue explants were incubated with Pil+Opa+ and Pil+ΔOpa GC (MOI~10) for 24 h and washed at 6 and 12 h to remove unassociated GC. Tissue explants were fixed, stained for ezrin, pMLC, GC and DNA, and analyzed using CFM. **(A, C)** Shown are representative images of ezrin **(A)** and pMLC **(C)** staining in both the luminal and subluminal layers of ectocervical epithelial cells. **(B, D)** The MFI of ezrin **(B)** and pMLC **(D)** in the luminal and the subluminal layers of the ectocervical epithelium. Data points represent individual randomly acquired images. **(E)** Shown are representative images of shedding cervical epithelial cells displaying ezrin accumulation at GC adherent sites (arrows). Scale bar, 10 μm. n=2. \*\*\**p*<0.001.

### GC induces the redistribution of Ezrin and NMII in endocervical epithelial cells, leading to F-actin reduction at GC adherent sites

We examined whether GC-induced F-actin reduction in endocervical epithelial cells was caused by ezrin and NMII reorganization, as what we observed in the polarized cell line model, using human cervical tissue explants. In the absence of GC, ezrin staining was concentrated at the apical surface of endocervical epithelial cells, colocalizing with apically polarized F-actin (Fig. 7A). After GC inoculation, ezrin staining was reduced exclusively underneath Pil+Opa+ and Pil+ΔOpa GC microcolonies at the apical surface of the endocervical epithelial cells, where F-actin reduction occurred (Fig. 7A, arrows). The FIR of ezrin staining underneath GC microcolonies relative to the adjacent no GC surface area was lower than 1 for both Pil+Opa+ and Pil+ΔOpa GC (Fig. 7B), suggesting ezrin disassociation from GC adherent sites. The FIR of ezrin staining at the apical surface relatively to the cytoplasm was also significantly reduced, suggesting ezrin disassociation from the apical surface to the cytoplasm (Fig. 7C). Both ezrin FIRs, the GC adherent site to the non-GC area and the apical to the cytoplasm, were significantly lower in Pil+ΔOpa GC than Pil+Opa+ GC colonized endocervical epithelial cells (Fig. 7B and 7C), suggesting that Opa expression inhibits ezrin redistribution induced by GC in the polarized endocervical epithelial cells. Thus, F-actin reduction at GC adherent sites of endocervical epithelial cells is concurrent with the localized disassociation of ezrin.

**Fig. 7.**
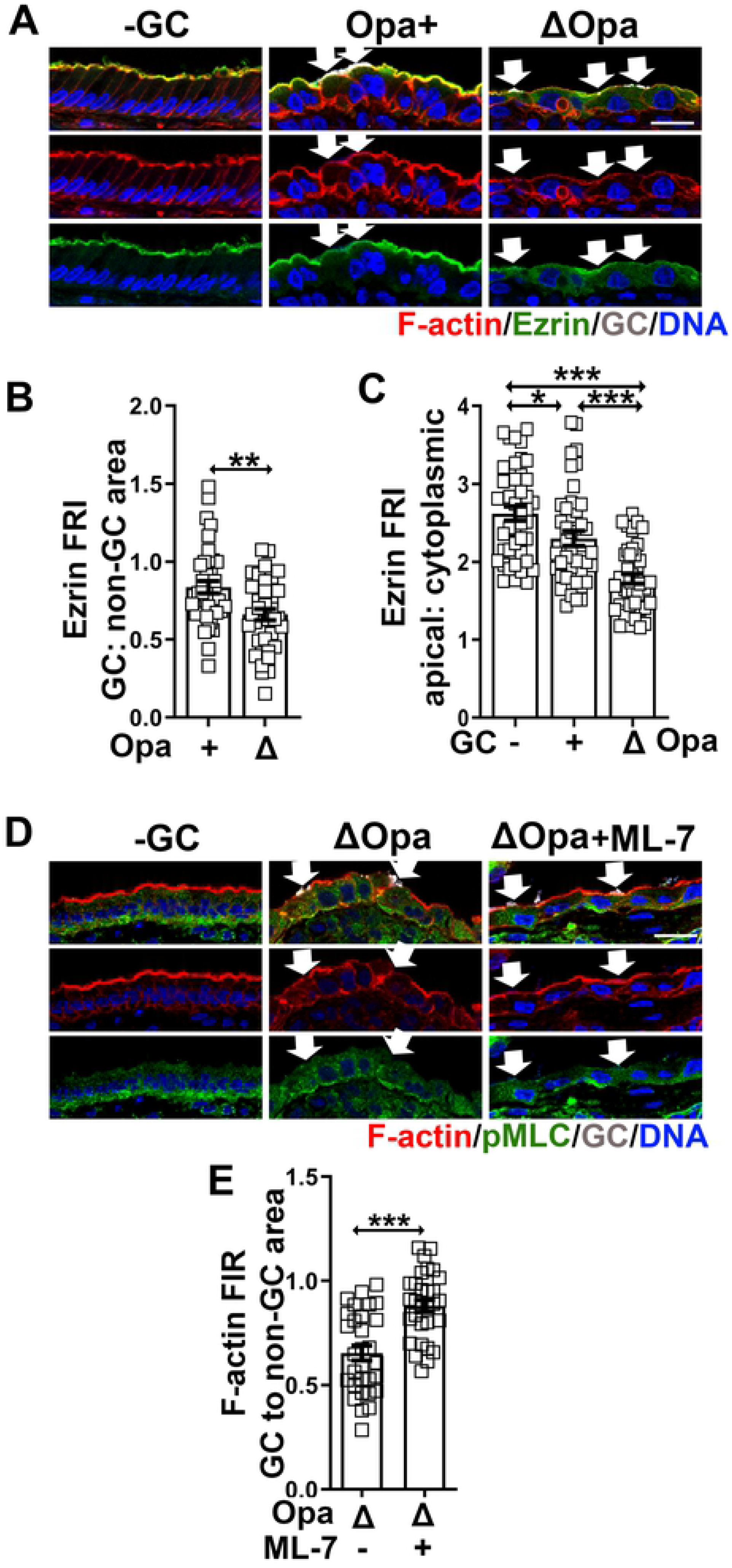
GC inoculation induces ezrin and NMII redistribution in the polarized endocervical epithelial cells, leading to F-actin reduction at GC adherent sites. Human cervical tissue explants were pretreated with or without ML-7 and incubated with Pil+Opa+ and Pil+ΔOpa GC (MOI~10) for 24 h in the absence or presence of ML-7 (10 μM). Tissue explants were stained for F-actin by phalloidin, ezrin, pMLC, GC and DNA, and analyzed using 3D-CFM. **(A, D)** Shown are representative images of ezrin **(A)** and F-actin **(D)** staining that intercept both the apical and basolateral surfaces of endocervical epithelila cells. Scale bar, 20 μm. **(B, C)** The redistribution of ezrin in the endocervical epithelium was quantified by the FIR (±SEM) of ezrin staining underneath individual GC microcolonies relative to the adjacent no GC surface area **(B)** and by the FIR at the apical surface relatively to the cytoplasm **(C)**. **(E)** The redistribution of F-actin was quantified by the FIR (±SEM) of phalloidin staining underneath individual GC microcolonies relative to the adjacent no GC area. Data points represent individual GC microcolonies. n=3. \*\**p*< 0.01; \*\*\**p*<0.001.

We have shown that GC inoculation induces NMII activation and recruitment to GC adherent site at the apical surface of the columnar endocervical epithelial cells [14]. Here, we determined whether such NMII activation led to F-actin reduction at GC adherent sites in endocervical epithelial cells. We pretreated human cervical tissue explants with or without the MLCK inhibitor ML-7 and incubated the explants with Pil+ΔOpa GC in the absence and presence of the inhibitor for 24 h. Tissue sections were analyzed using immunofluorescence microscopy, stained for GC, pMLC for activated NMII, F-actin by phalloidin, and DNA. We observed a recovery of F-actin staining underneath GC microcolonies in endocervical epithelial cells treated with ML-7, compared to those untreated ones (Fig. 7D, arrows). The FIR of phalloidin staining at GC adherent sites relative to no GC areas in the endocervical tissues treated with ML-7 (~0.88) was significantly increased compared to those untreated ones (~0.5) (Fig. 7E). These data suggest that GC-induced NMII activation is required for the F-actin reduction at GC adherent sites of the columnar endocervical epithelial cells.

## Discussion

While GC invasion into cervical epithelial cells has been well studied using both immortalized and primary cervical epithelial cells [17, 25, 26, 45, 52], whether GC could invade various types of cervical epithelial cells *in vivo* and how the heterogeneity of cervical epithelial cells impacted GC invasion was unknown. Using both cell line and human cervical tissue explant models, this study demonstrates that the polarity of epithelial cells and the expression of ezrin, an actin-plasma membrane linker, are two critical factors in epithelial cells that control GC invasion. GC prefer to invade into non-polarized over polarized epithelial cells, even though both were derived from the same cell line and express ezrin. GC prefer to invade into shedding cervical epithelial cells that are not polarized and express ezrin over non-polarized ectocervical epithelial cells that express no or little ezrin and polarized endocervical epithelial cells that express ezrin. GC invasion into epithelial cells requires the activation and recruitment of ezrin and ezrin-dependent actin accumulation at GC adherence sites. Reduced GC invasion into polarized epithelial cells is associated with drastic and localized decreases in ezrin and the actin cytoskeleton that are normally concentrated at the apical epithelial membrane. Our results reveal unique mechanisms utilized by GC to infect non-polarized squamous and polarized columnar epithelial cells.

Early studies have shown that both *Neisseria gonorrhoeae* [27, 35, 45, 53] and *Neisseria meningitidis* [37, 54, 55] induce the formation of cortical plaques, consisting of F-actin and ezrin, beneath adherent sites of their targeted host cells, epithelial and endothelial cells, respectively. The actin reorganization is associated with the elongation of host cell membrane protrusions surrounding bacteria, which strengthens bacterial adherence and favors the entry of the bacteria into host cells. Our study extends previous findings by providing strong evidence for an ezrin-dependent GC invasion mechanism. Here, we show that the inhibition of ezrin activation and recruitment in multiple ways, including an inhibitor, a decreased ezrin expression in the luminal layer of ectocervical epithelial cells, or GC-induced disassociation of ezrin from adherent sites in the apical membrane of polarized columnar endocervical epithelial cells, all inhibit GC invasion. Our results also provide mechanistic links between a series of events that leads to GC entry into epithelial cells. We show that GC interaction with non-polarized epithelial cells induces ezrin phosphorylation and recruitment to GC adherent sites. This ezrin activation is responsible for actin accumulation and microvilli elongation around GC microcolonies. In the absence of ezrin phosphorylation and recruitment, GC can no longer induce actin accumulation and microvilli elongation. These results show that activated ezrin links F-actin to the plasma membrane at GC adherent sites and promotes local actin polymerization, well-known functions of ezrin [32], which drives microvilli growth and promotes GC invasion. How GC adherence activates ezrin remains elusive. Pili has been shown to be required for the formation of the cortical plaque in epithelial cells [27, 35], and pili-mediated traction forces induce localized accumulation of F-actin and ezrin [34]. Our findings that Opa-expressing GC recruit more ezrin than those with all opa genes deleted and both Opa+ and ΔOpa GC increase ezrin phosphorylation suggest that Opa-host receptor interactions contribute to but not essential for the localized ezrin activation. GC adhesion has been shown to induce the recruitment of host receptors, such as CEACAMs [13] and epidermal growth receptors [35, 44, 52], which potentially activate and recruit ezrin through signaling and cell adhesion complexes.

The new findings of this study are that cell polarity is a critical host factor that curtails GC invasion into epithelial cells and that GC alter their infection mechanisms based on the polarity of epithelial cells. Even though ezrin and the actin cytoskeleton are highly enriched underneath the apical surface of columnar epithelial cells that GC directly interact with, GC induce sharp reductions rather than further enrichment of ezrin and F-actin and ablation rather than elongation of microvilli at adherent sites of polarized epithelial cells. Such surprising reorganization of ezrin, the actin cytoskeleton, and the apical morphology, opposite to the responses of squamous non-polarized epithelial cells, leads to a reduced GC invasion. Cell polarity is a primary factor distinguishing epithelial cells in the ecto- and endocervix. Evenly distributed cortical actin networks and ezrin in the squamous epithelial cells support a flat cell morphology and randomly appearing microvilli. In contrast, the highly polarized actin cytoskeleton and ezrin in columnar epithelial cells support their vertically extended morphology through the apical junction and dense microvilli at the apical membrane [30]. Such highly concentrated and organized actin networks under the dense apical microvilli make the apical membrane inflexible, preventing GC from expanding intimate interactions with epithelial cells. Our finding of GC-induced reduction of ezrin and F-actin at GC adherent sites suggests that GC overcome dense and rigid microvilli by disassembling the actin-ezrin framework of microvilli. Ablating the dense and rigid microvilli allows GC to expand their interaction with the epithelial cell membrane and secure colonization.

This study reveals that Ca^2+^-dependent activation of NMII is a mechanism by which GC trigger the disassembly of the apical actin networks in columnar epithelial cells, as both MLCK and Ca^2+^ flux inhibitors inhibit GC-induced reduction in F-actin. We further show that this mechanism is unique to polarized columnar epithelial cells, as GC do not appear to activate Ca^2+^ flux in non-polarized epithelial cells. Furthermore, such actin disassembly is localized and only occurs underneath the GC-epithelial membrane contact, suggesting that persistent and direct interactions between GC and epithelial cells are required. Our previously published data have shown that GC induce Ca^2+^-dependent activation of NMII in both polarized T84 cells and endocervical epithelial cells in tissue explants, which leads to the disassembly of the apical junction, allowing GC to transmigrate across the epithelium [14]. The actin cytoskeleton, organizing as a perijunctional actomyosin ring, is essential for the assembly, maintenance, and dynamic function of the apical junction [56]. GC-induced reorganization of the apical actin networks in polarized epithelial cells shown here is a potential cause of the apical junction disassembly. How GC activate Ca^2+^-flux exclusively in columnar epithelial cells remains an interesting question.

Our findings that Opa expression promotes GC invasion into T84 epithelial cells, no matter if they are non-polarized and polarized using gentamicin resistant assay, are consistent with previous reports [45]. Our TEM analysis also found a slightly higher percentage of intracellular bacteria in epithelial cells inoculated with Opa+ GC than ΔOpa GC, indicating that increases in gentamicin-resistant Opa+ GC result from invasion but not the differences in bacterial aggregation between Opa+ and ΔOpa GC [57]. Here we further show that Opa expression enhances ezrin and F-actin accumulation and microvilli elongation in non-polarized epithelial cells and inhibits ezrin-actin network disassembly and microvilli ablation in polarized epithelial cells. These results suggest that regulating the actin cytoskeleton is a mechanism by which Opa proteins facilitate GC invasion. As 10 out of 11 Opa proteins expressed by MS11 strains bind to CEACAMs [58], Opa-CEACAM interactions likely play the dominant role in regulating the epithelial actin cytoskeleton. CEACAMS are expressed on the luminal surface of both ecto- and endocervical epithelial cells, suggesting that such regulation potentially occurs *in vivo*. CEACAMs are known to interact with actin cytoskeleton proteins directly and indirectly, mediating cell-cell adhesion [59, 60], supporting that Opa proteins promote actin reorganization for GC invasion through binding to CEACAMs.

Surprisingly, we did not detect significant GC staining within the epithelia of both the ectocervix and endocervix, suggesting that GC invasion into epithelial cells probably is not a major pathway for GC infection of the human cervix, when compared to colonization and transmigration. The tissue explant model limits our experimental approaches to immunofluorescence microscopy, which is not as sensitive as the gentamicin resistant assay. However, we clearly visualized intracellular bacteria in epithelial cells shedding from the cervical epithelia using immunofluorescence microscopy, indicating that this approach is sensitive enough to detect invaded GC. Intracellular bacteria in shedding epithelial cells are consistent with previous observations using patients’ samples [51]. A lack of intracellular GC staining does not exclude the possibility of a small number of GC invaded into epithelial cells. The low levels of intraepithelial GC are associated with a lack of ezrin expression in ectocervical epithelial cells and highly polarized actin-ezrin networks in endocervical epithelial cells, which supports that ezrin is required for while highly polarized actin-ezrin networks inhibit GC invasion into epithelial cells. The luminal layer of ectocervical epithelial cells loses ezrin expression, likely due to cornification, a terminal differentiation process for turning over and renewing of the ectocervical epithelium. Here, we reveal that this cornification process also inhibits GC invasion into ectocervical epithelial cells.

This study sheds critical light on the mechanisms that both GC and human cervical epithelial cells develop to co-survive in the female reproductive tract. The reduced ezrin expression in the luminal layer of the ectocervical epithelial cells and the highly polarized actin-ezrin networks of endocervical epithelial cells protect the cervix from GC invasion. GC counteract by differentially modifying epithelial actin and microvilli organization, taking advantage of ezrin-actin networks when they are available and manipulatable, and disassembling ezrin-actin networks when they work against GC.

## Materials and Methods

### Neisseria strains

*N. gonorrhoeae* strain MS11 that expressed phase variable Pili and Opa (Pil+Opa+) and MS11 with all 11 *opa* gene deletion (Pil+ΔOpa) [61] were used. GC were grown on plates with GC media and 1% Kellogg’s supplement (GCK) for 16–18 h before inoculation. Pil+ colonies were identified based on colony morphology using a dissecting light microscope. GC was suspended, and the concentration of bacteria was determined by using a spectrophotometer. GC were inoculated at MOI ~10.

### Human epithelial cells

Human colorectal carcinoma cell line T84 cells (ATCC) was maintained in Dulbecco’s modified Eagle’s medium: Ham F12 (1:1) supplemented with 7% heat-inactivated FBS at 37°C with 5% CO_2_. To establish polarized epithelium, T84 cells were seeded at 6 ×10^4^ per transwell (6.5 mm diameter, Corning) and cultured for ~10 days until transepithelial electrical resistance (TEER) reached ~2000 Ω. TEER was measured using a Millicell ERS volt-ohm meter (Millipore). To establish non-polarized epithelium, cells were seeded at 1.2 ×10^5^ per transwell and cultured for 2 days. The 2- and 10-day cultures had the same number of T84 cells per transwell.

### Immunofluorescence analysis of epithelial cells on transwells

Non-polarized and polarized T84 cells were pre-treated with or without the ezrin inhibitor NSC668394 (20 μM, EMD Millipore), the myosin light chain kinase (MLCK) inhibitor ML-7 (10 μM, EMD Millipore), or the Ca^2+^ inhibitor 2APB (10 μM, EMD Millipore) for 1 h, and incubated with GC (MOI=10) for 6 h in the presence or absence of inhibitors. Cells were rinsed, fixed with 4% paraformaldehyde, permeabilized, and stained with phalloidin (Life Technology) and antibodies specific for ezrin (Santa Cruz Biotechnology), pMLC (Cell Signaling Technology) and GC, and Hoechst for nuclei (Life Technologies). Cells were analyzed by confocal fluorescence microscopy (Zeiss LSM 710, Carl Zeiss Microscopy LLC). Z-series of images were obtained in 0.57 μm slices from the top to the bottom of the monolayer, and three-dimensional (3D) composites were obtained using Zen software.

The redistribution of F-actin, ezrin, and pMLC induced by GC inoculation at the luminal surface was examined using xz images across the top to the bottom of epithelia. Fluorescence intensity (FI) profiles along a line underneath each GC microcolony and a line along the adjacent epithelial cell surface without GC were generated using NIH ImageJ software. The fluorescence intensity ratio (FIR) of F-actin, ezrin, and pMLC underneath GC microcolonies relative to the adjacent no GC epithelial surfaces was determined. A total of 54~81 microcolonies from three independent experiments were quantified.

### Human cervical tissue explants

Cervical tissue explants were cultured using a previously published protocol [12]. Briefly, cervical tissues were obtained from patients (28-40 years old) undergoing voluntary hysterectomies through National Disease Research Interchange (NDRI) and received within 24 h post-surgery. Muscle parts of the tissue were removed by using a carbon steel surgical blade. Remaining cervical tissues were cut into ~2.5 cm (L) × 0.6 cm (W) × 0.3 cm (H) pieces, incubated in the medium CMRL-1066 (GIBCO), containing 5% heat-inactivated fetal bovine serum, 2 mM L-glutamine, bovine insulin (1 μg/ml, Akron Biotech), and penicillin/streptomycin for 24 h and then in antibiotic-free media for another 24 h. Individual pieces of cervical tissue explants were pretreated with or without the MLCK inhibitor ML-7 (10 μM) and inoculated with GC at MOI ~10 (10 bacteria to 1 luminal epithelial cell) in the presence or absence of ML-7. The number of epithelial cells at the luminal surface was determined by the luminal surface area on individual pieces divided by the average luminal area of individual cervical epithelial cells (25 μm^2^). GC were collected from plates after ~18-h culture and resuspended in pre-warmed antibiotic-free cervical tissue culture media before inoculation. The concentration of bacteria was determined using a spectrophotometer. The cervical tissue explants were incubated with GC at 37°C with 5% CO_2_ with gentle shaking for 24 h. The infected cervical tissue explants were rinsed with antibiotic-free cervical tissue culture medium at 6 and 12 h to remove non-adhered bacteria.

### Immunofluorescence analysis of human cervical tissue explants

Tissue explants were fixed by 4% paraformaldehyde 24 h post-inoculation, embedded in 20% gelatin, cryopreserved, and sectioned by cryostat across the luminal and basal surfaces of the epithelium. Tissue sections were stained for F-actin by phalloidin (Cytoskeleton), ezrin (Santa Cruz Biotechnology), pMLC (Cell Signaling Technology), and GC by specific antibodies, and nuclei by Hoechst (Life Technologies), and analyzed using a confocal fluorescence microscope (Zeiss LSM 710, Carl Zeiss Microscopy LLC) as previously described [12]. Images of epithelia were randomly acquired as single images or Z-series of 0.57 μm/image, and 3D composites were generated using Zeiss Zen software.

GC distribution in cervical tissue explants was measured by the percentage of GC fluorescence intensity (% GC FI) at the luminal surface, within the cervical epithelium, and at the subepithelium. Seven randomly acquired images of each cervical region from each of three cervixes were quantified. The relative staining levels of ezrin or pMLC in the luminal layers and sub-luminal layers of ectocervical epithelial cells were evaluated by mean fluorescence intensity (MFI) in individual images. Six randomly acquired images from each of two cervixes were quantified. The redistribution of F-actin and ezrin in endocervical epithelial cells was evaluated by the fluorescence intensity ratio (FIR) underneath GC microcolonies relative to the adjacent surface areas without GC attached, or at the apical surface relative to in the cytoplasm in individual endocervical epithelial cells. A total of 27~52 microcolonies from images of three cervixes were quantified.

### GC adherence and invasion assays

The adherence and invasion of GC in T84 cells were performed as previously described [26, 44]. Briefly, non-polarized and polarized T84 cells were pretreated with or without ezrin inhibitor, NSC668394 (20 μM), for 1 h, inoculated with GC in the presence or absence of the inhibitor from the apical chamber for 3 h for analyzing adherence and 6 h for analyzing invasion. For adherence analysis, GC-inoculated cells were washed intensively, lysed, and plated on GCK plates to determine the CFU. For invasion analysis, GC-inoculated cells were treated with gentamicin (100 μg/ml) for 2 h, washed, lysed, and plated on GCK plates to numerate CFU. Gentamicin-resistant bacteria were counted as invaded GC. Three transwells from each of three independent experiments were quantified.

### Calcium imaging

Non-polarized and polarized T84 cells on transwells were pretreated with or without thapsigargin (10 μM, Sigma) for 1 h and incubated with GC (MOI~10) apically with or without the inhibitors for 4 h. Thapsigargin is an endoplasmic reticulum Ca^2+^ ATPase inhibitor that causes Ca^2+^ elevation in the cytoplasm and thereby served as a positive control. Cells were incubated with the fluorescent calcium indicator Fluo-4 (100 μM, Life Technologies) for 1 h, and xz images were acquired in the presence of the membrane dye CellMask (5 mg/ml, Life Technology) using Leica TCS SP5X confocal microscope (Leica Microsystems). The MFI of Fluo-4 in the cytoplasmic region in individual cells was measured using the NIH ImageJ software. Twenty randomly selected cells from each of three independent experiments were quantified.

### Function analysis of the apical junction

Polarized T84 cells were incubated with GC apically for 6 h. The CellMask dye (5 μg/ml, Life Technologies) was added in the basolateral chamber during the last 15 min, and xz images were acquired using Leica TCS SP5 X confocal microscope (Leica Microsystems). The percentage of epithelial cells displaying CellMask staining at the apical membrane in each randomly acquired image was determined.

### Western blotting

Non-polarized and polarized T84 cells were pretreated with or without the ezrin inhibitor NSC668394 (20 μM) for 1 h and incubated with GC with or without the inhibitor from the top chamber for 6 h, and then lysed by RIPA buffer [0.1% Triton x100, 0.5% deoxycholate, 0.1% SDS, 50 mM Tris-HCl, pH 7.4, 150 mM NaCl, 1 mM EGTA, 2 mM EDTA, 1 mM Na_3_VO_4_, 50 mM NaF, 10 mM Na_2_PO_4_, and proteinase inhibitor cocktail (Sigma-Aldrich, St. Louis, MO)]. Lysates were resolved using SDS-PAGE gels (BioRad) and analyzed by Western blot. Blots were stained for ezrin (Cell Signaling Technology) and phosphorylated ezrin at T567 (Abcam), stripped and reprobed with anti-β-tubulin antibody (Sigma). Blots were imaged using a Fujifilm LAS-3000 (Fujifilm Medical Systems). Each data point represents individual transwells. Two transwells from two to three independent experiments were quantified.

### Transmission electron microscopic analysis

Non-polarized and polarized T84 cells were pretreated with or without the ezrin inhibitor NSC668394 (20 μM) for 1 h, and incubated with GC with or without the inhibitor from the top chamber for 6 h with MOI ~50. The epithelial cells on the transwell membrane were fixed, processed, and embedded using EPON 812 resin (Araldite/Medcast; Ted Pella). Thin sections crossing the top and the bottom membranes were collected, stained with uranyl acetate and lead citrate, and imaged by a ZEISS 10CA electron microscope (ZEISS). Microvilli elongation and ablation at the luminal membrane were quantified using 3~4 randomly acquired images at a magnification of 5k from each of 3 independent experiments. The percentage of intracellular GC was quantified using two images showing intracellular GC from each of 3 independent experiments.

### Statistical analysis

Statistical significance was assessed by using Student’s *t*-test by Prism software (GraphPad). P-values were determined using unpaired *t*-test with Welch’s correction in comparison with no infection controls.

## Ethics statement

Human cervical tissue explants were obtained through the National Disease Research Interchange (NDRI, Philadelphia, PA). Human cervical tissues used were anonymized. The use of human tissues has been approved by the Institution Review Board of the University of Maryland.

## Acknowledgments

We thank the UMD CBMG Imaging Core for all microscopy experiments and the Laboratory for Biological Ultrastructure for all the transmitted electron microscope experiments.

